# Beyond associations: A benchmark Causal Relation Extraction Dataset (CRED) of disease-causing genes, its comparative evaluation, interpretation and application

**DOI:** 10.1101/2024.09.17.613424

**Authors:** Nency Bansal, R C Sri Dhinesh, Ayush Pathak, Manikandan Narayanan

**Affiliations:** Department of Computer Science and Engineering, Indian Institute of Technology (IIT) Madras, Chennai - 600036, India; Centre for Integrative Biology and Systems Medicine, IIT Madras, Chennai - 600036, India; Department of Biotechnology, Alagappa College of Technology, Anna University, Chennai - 600036, India; Department of Computer Science and Engineering, Indian Institute of Information Technology (IIIT) Vadodara, Gandhinagar - 382028, India; Wadhwani School of Data Science and AI, IIT Madras, Chennai - 600036, India

## Abstract

Information on causal relationships is essential to many sciences, including biomedical science, and beneficial (e.g., causative rather than merely associative gene-disease relations can lead to better treatments). Despite much work on Relation Extraction (RE), automatically extracting causal relations from large text corpora remains less explored. Few existing studies on CRE (Causal RE) are limited to extracting causality within a sentence or for a particular disease, mainly due to the lack of a diverse benchmark dataset. Here, we carefully curate a new CRE Dataset (CRED) of 3639 (causal and non-causal) gene-disease pairs, spanning 204 diseases and 500 genes, within or across sentences of 267 published abstracts. CRED is assembled in two phases to reduce class imbalance, and its inter-annotator agreement is 89%. To assess CRED’s utility in classifying causal vs. non-causal pairs, we compared multiple classifiers and found SVM (Support Vector Machine) trained on embeddings from a deep learning transformer model called BioBERT to perform the best (F1 score 0.70). CRED outperformed a state-of-the-art RE dataset in terms of classifier performance and model interpretability, i.e., whether the model focuses importance/attention on words with causal connotations in abstracts. Moving from benchmark to real-world settings, application of our CRED-trained BioBERT+SVM model on all PubMed abstracts on Parkinson’s disease (PD) revealed both well- and less-studied PD-causing genes. For instance, genes predicted to be causal for PD in at least 50 abstracts by our model were already linked to PD in books; and lends confidence to further explore the other genes predicted to be causal in fewer abstracts. Our systematically curated and evaluated CRED, and its associated classification model and gene-disease causality scores, thus offer concrete resources for advancing future research in CRE from biomedical literature.

## Introduction

A knowledge of cause-effect relations is crucial in many fields, including biomedicine where the nature of relationship between entities like genes and diseases can influence treatment strategies (i.e., targeting genes that cause rather than respond/correlate to a disease can lead to more effective treatments); economic/social sciences where causal explanations are frequently sought for historical/forecasted events; machine-learning (ML) where side-information on causal vs. correlative features of a ML model can help improve its generalizability and interpretability^1^, and to avoid misleading conclusions (such as asthma being suggested as a preventive measure against mortality due to the confounding factor of asthmatics receiving more intensive therapy^2^); and finally natural language processing (NLP) where causal knowledge can improve text summarization and question-answering tasks^3^. However, manually extracting causal relations from vast corpora (e.g., gene-disease relations from the biomedical corpora PubMed with approximately 35 million articles) is impractical; and advanced NLP methods, being increasingly used for automated extraction and summarization of information from literature, need to be extended to perform causal text mining.

Extracting useful information from published articles, specifically RE, is a mature and well-established subfield of NLP. In contrast, the area of CRE from text is in a nascent stage, mainly due to the lack of ground-truth datasets. For instance, focusing on gene-disease relations in biomedical literature, there are more RE than CRE datasets (see summary in Table 1, and details of these related works in Section A.1 of Appendix). A pioneering CRE dataset and associated model pertains to the Sjögren’s Syndrome (SS)^4^, but it is limited to extracting causal factors of only one disease (SS) and can consider only a single sentence at a time. Extending such datasets to capture information on causal relations between more genes and diseases, spread across multiple sentences, is important to understand disease mechanisms and advance drug discovery. It is also challenging and time-consuming to extract causal rather than associative gene-disease relations from the PubMed corpus, due to the scarcity and ambiguity of mentions of causal relations in research articles. It is also important to note that RE databases catalog associative links, often mixing mechanistic claims (“Mutation in Gene X causes Disease Y”) with co-occurrences. This mixing hampers models designed to distinguish true causality from mere correlation, and underscores the need for datasets that are explicitly built for the purpose of CRE.

**Table 1.** Information on existing RE and CRE datasets. A subset of the total pairs (Tot Pairs) are positive relations (Pos Rel), with positive class referring to causal relationship for CRE and any (including correlative) relationship for RE datasets. Categorizing a dataset as RE vs. CRE is based on the cited publication describing the dataset. Drugs are also viewed as chemicals in EU-ADR and ADE datasets. Targets may also contain genes in EU-ADR dataset. Abbreviations: D - Disease, G - Gene, C - Chemical, Dr - Drug, T - Target and V - Variant; # - “Number of” (a number that we couldn’t find in the paper or website is marked as NA); aka - also known as. Other abbreviations such as names of datasets are expanded in Section A.1 of Appendix.

| Dataset | Entity Type | CRE/RE | # Tot Pairs | # Pos Rel | Example text from dataset |
| --- | --- | --- | --- | --- | --- |
| SemEval<sup>7</sup> | General Relations | CRE | 10717 | 1331 | *The burst has been caused by water hammer.* |
| ADE<sup>8</sup> | Dr→Effect | CRE | 23516 | 6821 | *The patient developed a rash after taking penicillin.* |
| Sjögren’s Syndrome<sup>4</sup> | Factor→SS | CRE | 383 | 383 | *Hypokalemic paralysis is a rare presentation of Fanconi syndrome (FS) caused by Sjogren’s Syndrome (SS).* |
| GAD<sup>9</sup> | G-D | RE | 5330 | 2756 | *The patient had a mutation in the BRCA1 gene, which was associated with an increased risk of breast cancer.* |
| EU-ADR<sup>10</sup> | Dr-D, T-Dr, T-D | RE | 355 | 261 | *The patient was taking methotrexate for rheumatoid arthritis and developed pneumonitis.* |
| CDR<sup>11</sup> (aka BC5CDR) | C-D | RE | 12850 | 3116 | *Intravenous administration of lidocaine in a 67-year-old man resulted in profound depression of the sinoatrial nodal pacemakers.* |
| BioRED<sup>12</sup> | D-G, D-C, D-V, C-C, C-G, G-G, V-C, V-V | RE | NA | 6503 | *A polymorphism of C-to-T substitution at -31 IL1B is associated with the risk of advanced gastric adenocarcinoma in a Japanese population.* |

To address the above-mentioned gaps in CRE research, our study (Figure 1a) makes these contributions:

**Figure 1.**
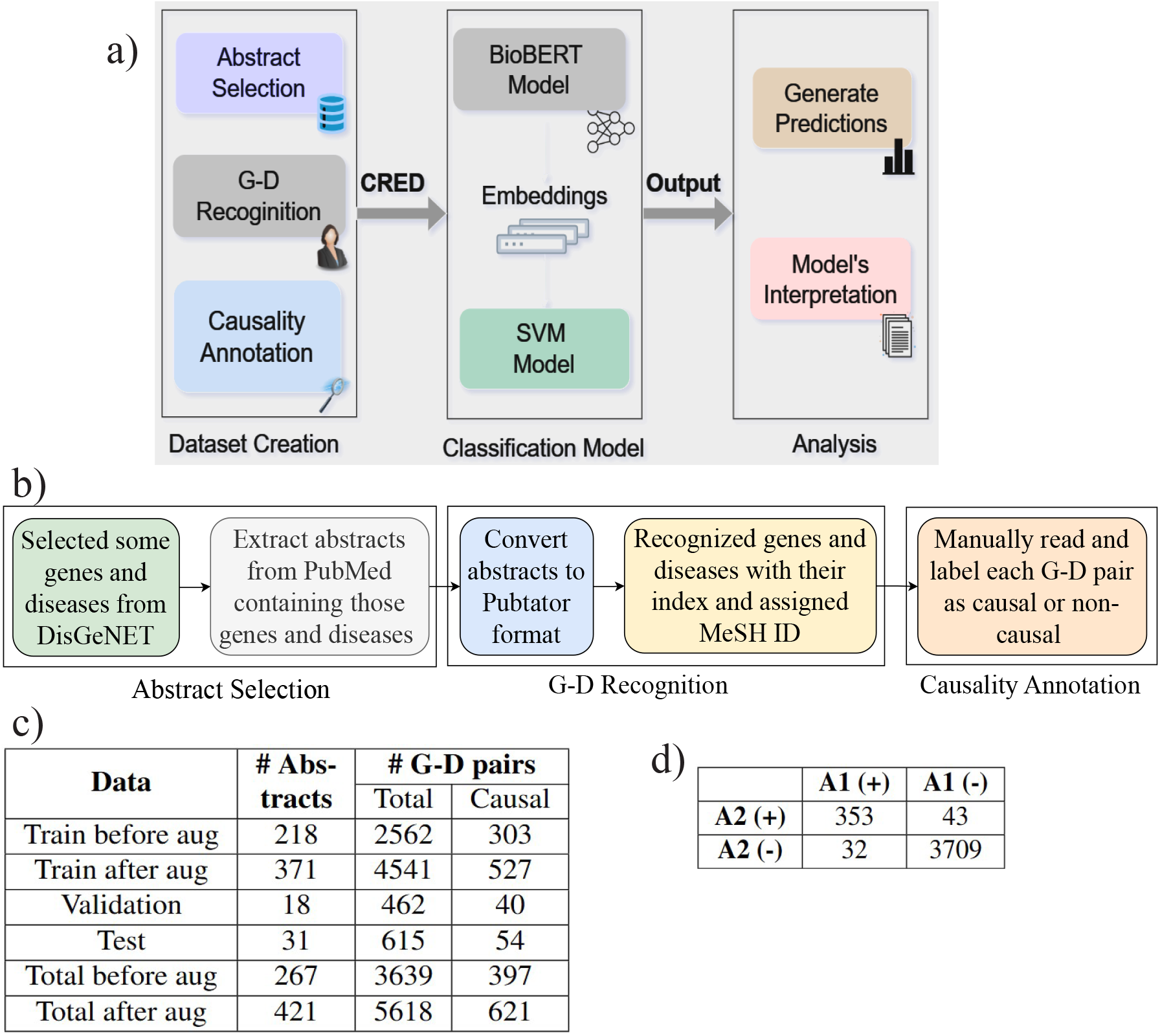
(a) Overview of our work, (b) Steps involved in CRED creation (Phase 1), (c) CRED Statistics. # represents the number, (d) Contingency matrix. A1 and A2 refer to two different annotators. (+) and (−) represents causal and non-causal classes respectively.

1. We have created CRED, a dataset of disease-causing genes extracted from published biomedical abstracts in PubMed^5^. It is a first-of-its-kind dataset that includes both intra- and inter-sentence gene-disease (G-D) pairs, spanning a diverse set of 500 genes and 204 diseases. CRED exhibits an inter-annotator agreement of 0.89 (Cohen’s kappa) as a result of thoughtful manual curation guidelines.
2. To assess the utility of our CRED, we have trained or fine-tuned different state-of-the-art (SOTA) classifiers, including two Large Language Models (LLMs), on CRED. We found that a SVM model trained on BioBERT^6^ embeddings performs best with an F1 score of 0.70 to distinguish causal from non-causal G-D pairs.
3. We interpret the predictions of our CRED-trained classification model and compare it with model interpretation scores of a RE dataset trained model to verify if the CRED model actually focuses on the contexts or words with causal connotations (causal-connotation words) to make correct predictions. Our model indeed pays attention to such causal words and has learned the context well without focusing on the names of specific genes and diseases.
4. In a real-world application of our CRED-trained model to all abstracts mentioning Parkinson’s Disease (PD), our model could extract known/well-studied PD-causing genes, as well as quantify the evidence of causality of numerous less-studied genes. In another application, CRED-wide gene-disease causality scores revealed common genes that cause multiple diseases.

These contributions, including a gold-standard database of manually curated causal relations and results from extensive evaluations of models trained using our dataset (to learn the fine-grained linguistic signals underlying a cause-effect relation), show that reliable CRE from biomedical literature is possible.

## Methods

This section describes the steps involved in creating our CRE dataset CRED, assessing its utility to perform classification-based causal text mining, and interpreting the classification model’s predictions. When creating CRED, we assume a “disease-causative” gene to be defined as a gene that upon perturbation (including over-expression, knockout, or any of its mutations) changes the status or severity of the disease (e.g., leads to manifestation/activation, suppression, or altered severity of the disease).

### CRED Creation

A main contribution of our work is in creating a CRED dataset of 5618 causal and non-causal G-D pairs (Figure 1c), including 1722 intra-sentence and 3896 inter-sentence G-D pairs, which are systematically extracted and annotated based on 267 abstracts in two phases as explained next.

#### Abstract Selection

In phase 1, to select abstracts, we start with a list of gene-disease pairs from DisGeNET repository^13^ with a Gene-Disease Association (GDA) score of at least 0.5 for causal relation and some random gene-disease pairs for non-causal relations. Second, we extract the abstracts mentioning the gene-disease pair from PubMed and restrict our attention to the top 10 abstracts (in terms of search relevance) for each gene-disease pair. Note that each abstract is identified or represented using its PMID (PubMed ID). Overall, we select 267 abstracts comprising a total of 3553 (including causal and non-causal) gene-disease pairs, spanning 204 diseases and 500 genes.

#### Gene-Disease Recoginition

After selecting the abstracts, we use PubTator^14,15^ to perform NER of gene and disease entities in the abstract and to group different representations of the same entity to a common ID called a MesH (Medical Subject Heading) ID for diseases and Entrez ID for genes. For example, the different representations of Parkinson’s disease, such as PD, Parkinson, or Parkinson’s, are all mapped to the same MesH ID D010300.

#### Causality Annotation

##### Task Overview

We followed a double-annotation approach, with the annotations of the first annotator used to prepare the CRED dataset and those of the second annotator used to assess Inter-Annotator Agreement (see IAA section below). Each annotator manually read each selected abstract and annotated each gene-disease pair as causal (1) or non-causal (−1) based on the text. The major problem was that very few abstracts in biology talk precisely about disease-causing genes; associations are more commonly reported in abstracts than causal relations. For example, an abstract mentioning two genes and five diseases may report only one of these ten gene-disease pairs as causal; the other nine pairs are given the non-causal label in our dataset. Figure 1b shows the steps involved in annotating the dataset. Figure 2a shows an easy example, and Figure 2b shows a complex example illustrating the CRED creation steps.

**Figure 2.**
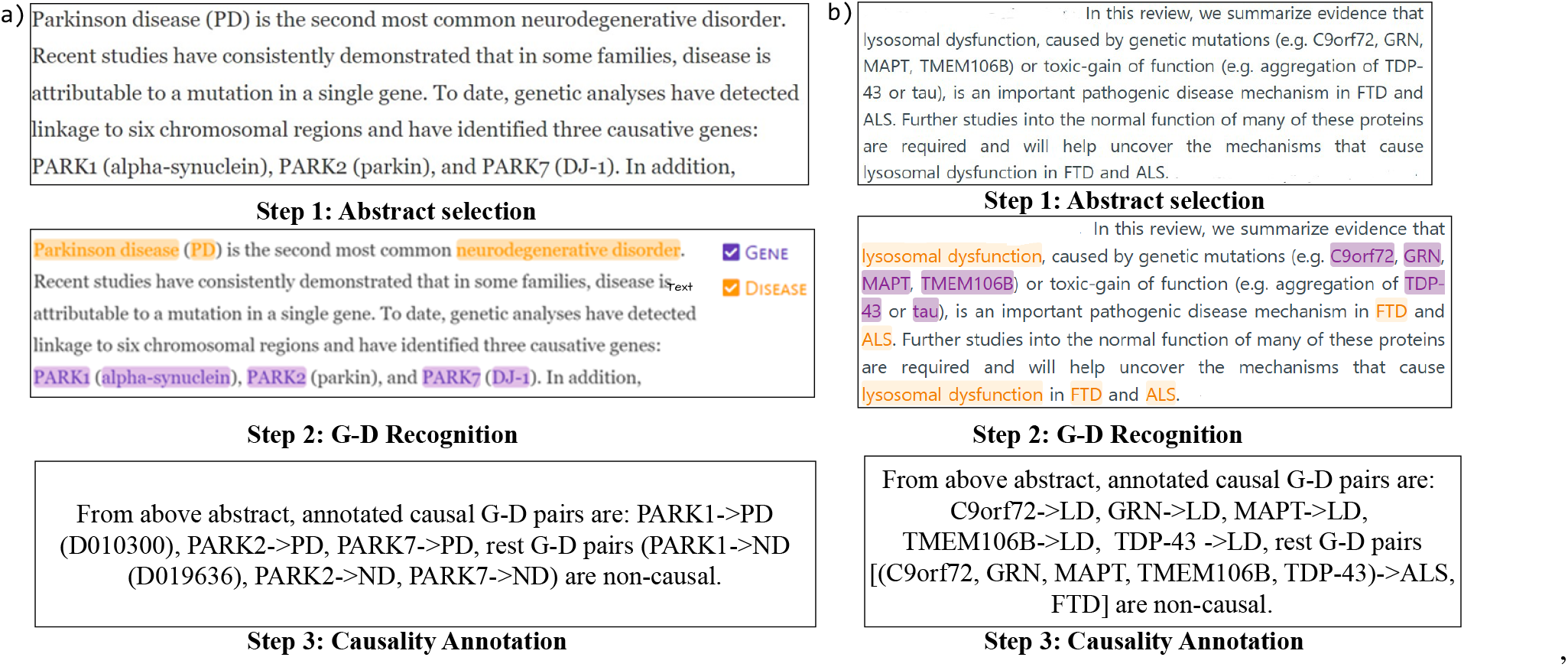
a) Simple example (PMID: 15717024) to understand the steps of CRED creation. ND denotes neurodegenerative disorder. b) A complex example (PMID:33812000). LD represents Lysosomal dysfunction. For more example, refer to Figure S1 in the Appendix.

##### Annotation guidelines

Annotation guidelines were formed under the guidance of a senior annotator. Following are the final annotation guidelines followed by the annotators:

1. The relation should be explicitly mentioned in the abstract and should not depend on pre-acquired knowledge of the annotator about the disease or gene.
2. The annotations must be performed both within as well as across sentences. The context (the rest of the abstract) can be used for disambiguation.
3. Association between genes and disease should not be considered as causal relations. Risk factors and therapeutic targets are not considered causal factors. Causality must be mentioned specifically.
4. Mention of causal relation does not only mean the presence of words like causing, leads to, etc., but also considers the context of the abstract or some biomedical terms like upregulation, perturbation, etc.
5. The gene-disease pair should be labeled as causal (Label=1) if the abstract clearly states that there is a causal relation between genes and diseases and all other relations (association, co-occurrence, disease activating gene, gene activating other genes) are considered as non-causal (Label=-1).
6. Annotators have to check if the recognition of entities by PubTator is correct. If not, then correct the incorrect/unidentified genes and diseases.

#### Phase 2 of creation of dataset

To address the class imbalance (few causal compared to non-causal pairs) in the dataset assembled in phase 1, we performed another phase of annotation. In phase 2, we selected 1000 new abstracts from PubMed which did not overlap with phase 1 (using the abstract selection process explained above), and passed all the G-D pairs (10,283) through the model (BioBERT+SVM) trained on phase 1 data. This model predicted 735 G-D pairs as causal, which were then provided to both annotators for manual annotation. During this manual annotation step, annotators also discarded all the missing and erroneous entities of PubTator. The causal annotations of the first annotator (specifically 191 causal pairs out of the 735) were added to CRED’s training data to reduce the class imbalance and those of the second annotator were used to assess IAA, following a similar process as in phase 1. Validation data and test data from phase 1 were not altered during phase 2, in order to prevent data leak.

#### Annotators and Inter-Annotator Agreement (IAA)

We followed a double-annotation approach, wherein two independent in-house annotators (co-authors of this article, referred to as A1 and A2) provided annotations to curate the causal relationship in each article in the combined phase 1 and 2 dataset. Annotator A1 is doing graduate-level research in the bioinformatics area, and A2 has completed his undergraduate degree (B.Tech) in the biotechnology stream. Hence, both annotators have the required amount of domain-specific knowledge. We used the annotations of A1 to prepare the CRED dataset and the annotations of A1 and A2 to evaluate IAA. The above-mentioned annotation guidelines were followed by both annotators to maintain consistency.

To assess the consistency of annotation among the annotators, we measured the agreement of annotations using Cohen’s kappa^16^. The contingency matrix in Figure 1d shows the number of agreements and disagreements. We achieved an inter-annotator agreement score of 0.893. Figure S2, S3 in Appendix show examples of discrepancy in annotations between the annotators.

### Classification Model

To assess the utility of our CRED dataset, we want to use it to train a CRE model that can predict if a given G-D pair in an abstract is causal or not.

#### Data Augmentation

During phase 1 alone, we augmented abstracts containing causal relations using techniques like “SynonymAug” from the “nlpaug” library^17^. This involved synonym replacement and spelling variations, ensuring the context remained unchanged, and was applied only to the training data. An example of an original abstract and its augmented version is shown in Figure S4 of Appendix. Augmentation in general helps address the class imbalance by letting the model train well from both positive and negative classes. In our case, we would like to use our augmented training data to improve upon the non-augmented model performance (reported in Table S1 of the Appendix). It is important to note that validation and test data were not augmented. Statistics on augmented data points is given in Figure 1c. Unless otherwise specified, CRED training data refers to the training data after augmentation. So all the analyses in this work performed on CRED training data use both the original and augmented data points.

#### Data Cleaning and Preprocessing

To enhance our model’s generalizability across various diseases and genes, we use data-cleaning techniques to remove stopwords. We substitute the disease name with @DiseaseTgt$ and gene name with @GeneSrc$. This approach focuses the model on the linguistic context and relationships rather than specific gene or disease names to help reduce bias towards well-studied genes/diseases and improve performance on unseen genes/diseases.

#### Classifiers

We tried different types of models for the classification of G-D pairs as causal or non-causal. These models can be grouped as:

1. BERT-based Model: We tried different BERT^18^ based models for comparison (such as BioBERT^6^, PubMedBERT^19^, SciBERT^20^, and BlueBERT^21^). We fine-tuned existing models on the CRED training dataset and tested them on the CRED test dataset to enable a fair comparison.
2. BERT-based Model with a (Downstream) Classifier: We also tried combining a BERT-based model with a downstream classifier model like SVM, XGBoost, or Random Forest. In a model of this type, preprocessed and cleaned text is passed as input to a transformer model such as BioBERT^6^; and contextualized embeddings generated from the transformer model are passed as input to train and test the downstream classifier such as SVM. We selected the encoded (contextualized) embeddings of only the gene and disease mentions in the abstract, rather than [CLS] token (special token containing the weighted average of all the tokens, generally used for classification tasks) embeddings. We concatenated the two selected embeddings, and passed them as input to the downstream classifier. We have taken the average of embeddings in an abstract in case of multiple occurrences of the same gene and disease. For details related to dimensions of embeddings, refer Section A.2 of Appendix. Note that among the different models of this type that we tested, BioBERT+SVM classifier performed the best (see Table 2) and hence used for other analyses (and referred to as “our model” in this work).
3. Large Language Model (LLM): Many LLMs are available, but we chose two open-source LLMs to fine-tune them and check their performance for the classification task. We selected MMed-Llama-3 (8 billion parameters) as it is a domain-specific LLM fine-tuned on biomedical corpora from PubMed Central^22^, and is also verified as a model performing well for biomedical natural language processing (BioNLP) tasks, including relation extraction^23^. We also selected Phi-4 (14 billion parameters) developed by Microsoft, as it is a recent high-performing LLM trained on high-quality publicly available data, educational data, synthetic “textbook-like” data created for the purpose of learning math, coding, common sense reasoning, and general knowledge of the world^24^.

**Table 2.**
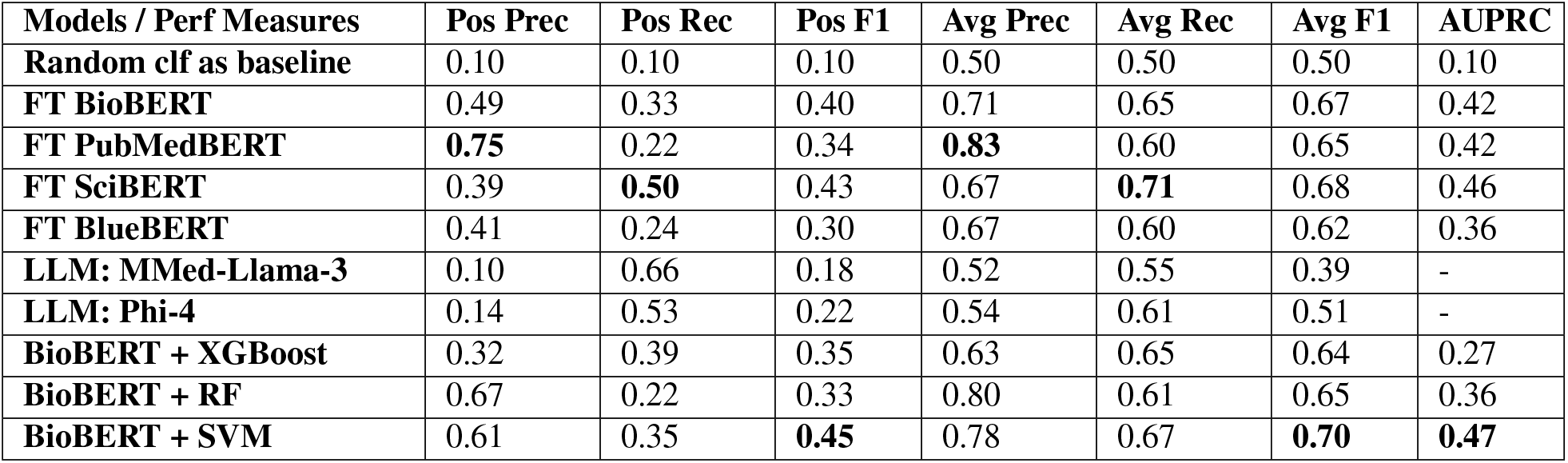
Performance of different SOTA models on CRED. Pos represents the causal class, whereas averages show the average performance of both classes. AUPRC of LLMs can’t be calculated as LLMs were trained only for classification. The “clf” represents the classifier. “FT” represents fine-tuned. “RF” represents Random Forest. “Pos” denotes the metrics for causal class and “Avg” contains scores averaged on both classes. Prec, Rec, and F1 denote precision, recall, and F1 score, respectively.

For fine-tuning LLMs, we use Group Relative Policy Optimization (GRPO)^25^, which is a reinforcement learning technique that can be efficiently implemented via updates of only low-rank adaptation (LoRA) matrices^26^. In more detail, we first provide the required format/schema of the prompt and response (e.g., prompt should be “Is @GeneSrc$ causal for @DiseaseTgt$ in the given text?” and response should be a binary Yes/No) and use the CRED training set as the training data to fine-tune the model. We next design a reward function required for GRPO by setting a reward of 10% for the correct schema, 70% for the correct response, and 20% for the partially correct response; and a penalty of 70% for the incorrect response.

#### Implementation Details

For the BERT-based and BERT-based with classifier models, we used an Amazon Web Services Elastic Computing Cloud (AWS EC2) instance running a 64-bit Ubuntu operating system on a configuration of 1 NVIDIA GPU with 24 GB memory, 32 vCPUs (Virtual CPUs), and 220 GB externally attached memory. Specification details of all the packages used for this work are mentioned in Table S2 of the Appendix section. It took around 20-25 minutes for models like BioBERT with SVM to train and test on CRED.

The compute resources mentioned above were not sufficient for fine-tuning LLMs, so we used an NVIDIA RTX 6000 Ada machine (GPU with 48 GB memory, 128 cores, and a 512 GB hard disk) for fine-tuning Phi-4, and it took around 12 hours. We used an NVIDIA A100 machine (GPU with 80 GB memory, 96 vCPUs, and 300 GB externally attached memory) for fine-tuning MMed-Llama-3, and it took around 40 hours.

#### Hyperparameter tuning

For each model of interest, the values of the hyperparameters tried, their corresponding performance on the validation data, and the hyperparameter configuration yielding the best validation performance are all mentioned in appropriate Supplementary Figures (e.g., Figure S5, S6 in the Appendix). The plots in these hyperparameter figures have been generated using the “Weights and Biases” platform^27^. Note that for each model, we select the hyperparameter with the best validation performance and apply the resulting model on unseen test data. This test performance is reported in the Results section.

### Model Interpretation

Interpreting our model’s predictions involves analysing the reasons behind the model’s classification. To do so, we calculate the importance score of each input token (denoted *w*) in a given abstract (denoted *Abstract*) towards an output prediction (for a given *G* − *D* pair in the *Abstract*). We first replace *w* by a special token “MASK” in the *Abstract* and apply the classifier to get the predicted probability for the *G* − *D* pair to be causal (denoted as the perturbed probability *p*_*pert*_). The corresponding predicted probability when the classifier is applied on the original abstract without masking/perturbation is denoted *p*_*orig*_. We then derive the importance score (*ImpScore*) as per Equation 1, inspired by another published feature importance score^28^.

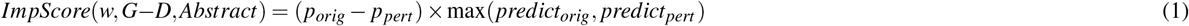

Here, *predict*_*orig*_ denotes the prediction of the model applied on the original abstract (i.e., 1 if the *G* − *D* pair is predicted to be causal and 0 otherwise; for probabilistic model based classifiers, *predict*_*orig*_ =⌊*p*_*orig*_ + 0.5⌋). Similarly, *predict*_*pert*_ denotes the model prediction on the perturbed/masked abstract. The higher the ImpScore of a token (*w*), the more that token is responsible for the prediction; it thus reveals the words of the input sentence focused by the model for making the prediction. A negative ImpScore shows that the word supports making the wrong prediction (since we only calculate these scores for ground-truth causal G-D pairs). Please refer Section A.3 of Appendix for more information on ImpScore.

### Applications of CRED - Methodological Details

We consider two applications of CRED: (i) predicting the causal genes of PD using all relevant papers in PubMed, and (ii) deriving a CRED-wide causality score of a G-D pair. The methodological details of these applications are in Section A.4 of Appendix.

## Results

To assess the utility of our CRE dataset for causal text mining, both in absolute terms and also relative to a SOTA RE dataset CDR, we train and test multiple classifiers on CRED and CDR and present these results in this section. Note that CDR (Chemical Disease Relation) dataset refers to the BC5CDR (BioCreative V CDR) corpus^11^. We report performance using precision, recall, and F1-score instead of accuracy to account for class imbalance. We also calculate the area under precision-recall curve (AUPRC) to analyze the performance of SOTA models. We divide our benchmark CRED into training, validation, and testing data, as shown in Table 1c. Considering the moderate size of CRED, we apply four-fold cross-validation on training+validation data, in which three folds are used for training, and one fold is used for validation. The validation fold is used for hyperparameter tuning, provided in Figure S5, S6 in the Appendix.

### Comparison with SOTA Methods

Table 2 compares different SOTA methods on CRED. Kindly refer Section A.2 and Table S3 in Appendix for a detailed explanation of the fine-tuning process and number of parameters of our model. All the values of hyperparameters tried, best values obtained, and associated performance results are shown in Figure S7, S8, S9, S10, S11, S12 in Appendix. This comparison (Table 2) showed that our BioBERT+SVM model has a good balance of precision and recall and hence the highest F1-score for the positive class as well as the average of both classes. Our model also outperforms all other SOTA models in terms of AUPRC.

We also compare our model’s performance with LLMs, MMed-Llama-3 and Phi-4. For a fair comparison, these models were fine-tuned on CRED training data before evaluating on the CRED test data. Our model outperforms these LLMs, probably due to a high number of parameters in the LLMs (in the billions) which makes them difficult to fine-tune using our dataset of 4541 training data points.

### Comparison with other datasets

To assess the utility of CRED, we compared our CRE dataset (CRED) with other established predominantly-RE datasets like CDR, BioRED and GAD, specifically to check if these datasets could be used to train a CRE model. For this purpose, we trained the BioBERT+SVM model on the CDR, BioRED, or GAD dataset and tested it on the CRED test data. Table 3 shows that this strategy leads to a significant drop in performance (compared to the Positive F1 score of 0.45 for the CRED-trained model), implying that the models trained using these other datasets are unable to identify causal relations from abstracts in CRED. This verifies the need for and supports our efforts to create an explicit CRE dataset. For completeness, Table S4 reports out-of-domain results of CRED-trained model tested on CDR, BioRED, or GAD.

**Table 3.** Performance of BioBERT+SVM model. Tr-Te represents training data - test data. So, CDR-CRED indicates that the model is trained on CDR dataset and tested on CRED. Prec, Rec, and F1 denote precision, recall, and F1 score respectively.

| Tr-Te | Positive class |  |  | Average |  |  |
| --- | --- | --- | --- | --- | --- | --- |
|  | Prec | Rec | F1 | Prec | Rec | F1 |
| **CDR-CRED** | 0.18 | 0.72 | 0.29 | 0.57 | 0.71 | 0.55 |
| **BioRED-CRED** | 0.06 | 0.01 | 0.01 | 0.48 | 0.50 | 0.47 |
| **GAD-CRED** | 0.10 | 0.69 | 0.17 | 0.51 | 0.54 | 0.36 |

### Experiment to assess model robustness

With BioBERT+SVM as the best-performing model, we next assessed the robustness of this model to different training datasets under a four-fold cross-validation framework shown in Figure S13. We indeed find the model to be robust.

### Experiment with different embeddings

We tried changing the input embeddings of our model, from the default “only G-D embeddings” to the “CLS token with G-D embeddings” and the “only CLS token embeddings”. The performance dipped with CLS and G-D embeddings and was poor with only CLS embeddings (see Figure 3). This performance trend is likely because CLS token’s embedding contains the weighted average of the complete abstract, which might not be sufficient to distinguish between causal and non-causal G-D pairs in the same abstract.

**Figure 3.**
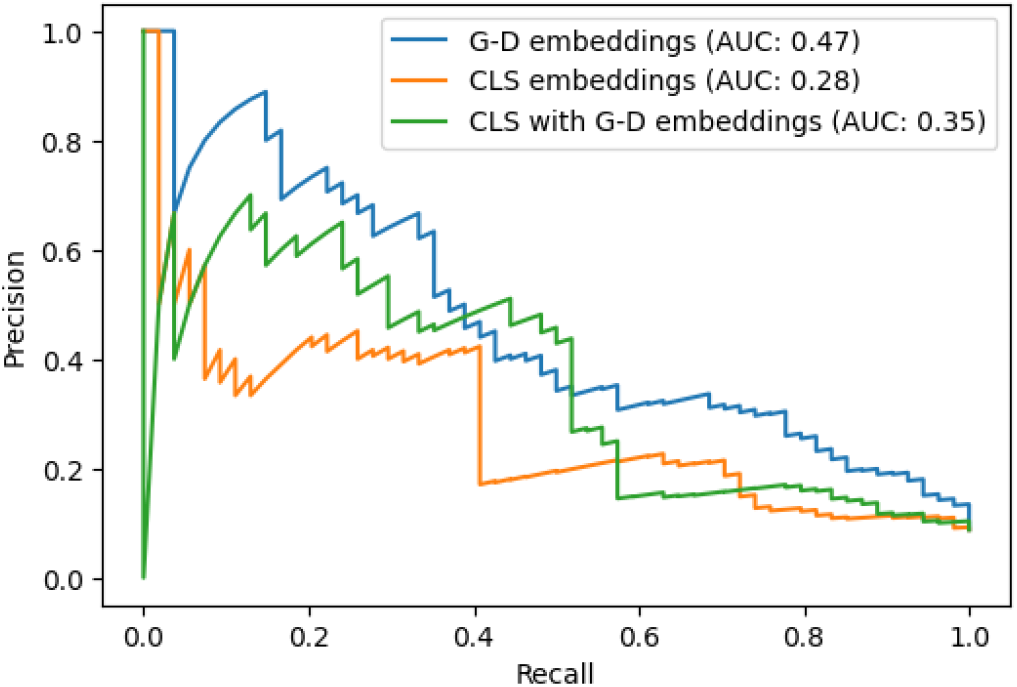
BioBERT+SVM performance when trained on CLS embeddings, CLS with G-D embeddings and only G-D embeddings. AUC represents area under the curve.

### Interpreting model predictions using importance scores

To assess the difference between models trained using CRE dataset (CRED) vs. RE dataset (CDR) and to interpret our model’s predictions, we identify the tokens/words *w*, a given model focuses on to make the predictions. Towards this end, we specifically use the importance score of (*w, Abstract*) tuples (see Methods). We would like to test if CRED-trained model assigns higher importance scores to causal-connotation words than association-related words, compared to the CDR-trained model. Towards this analysis, we selected synonyms of the words “cause” and “associate” (mentioned in section A.5 of the Appendix). For a matching word (or synonym) in each abstract, we take the maximum importance score in case of multiple occurrences of the word. In detail, we take the maximum importance score of all the matching words for (*w, Abstract*) tuples in true positives of CRED train and test data separately.

Figure 4a,b shows the importance scores calculated using the CRED-trained BioBERT+SVM model. From this figure, we observe that the importance scores of causal-connotation words are significantly higher than that of associate-related words in the training data (Wilcoxon one-sided test P-value = 0.00003). Our CRED-trained BioBERT+SVM model thus focuses more on causal-connotation words as compared to associate-related words across all true positives in the train data. A similar trend is also seen in the test data (Wilcoxon one-sided test P-value = 0.01180), with the marginal significance of the P-value likely due to the smaller number of matching words for (*w, Abstract*) tuples in the test data. To provide additional context for these results, we repeated this importance score analysis using other models with poorer classification performance like the CRED-trained BioBERT+XGBoost model, and out-of-domain-trained models like the CDR-trained BioBERT with SVM and XGBoost models, and present and interpret these results in Figures S14, S15 and S16 in Appendix.

**Figure 4.**
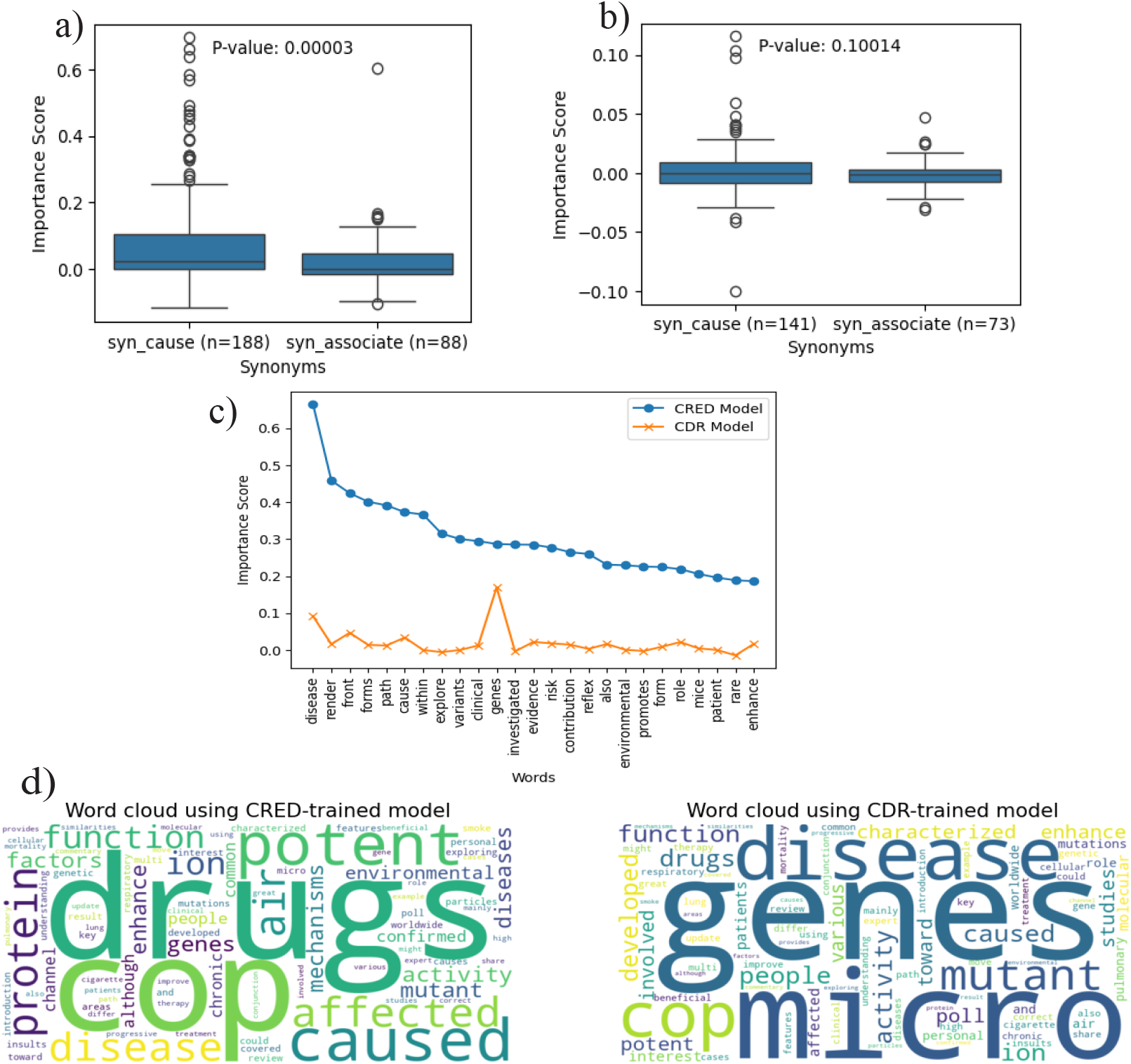
Interpretation Results. (a,b) Comparison of importance scores of “cause” vs. “associate” synonyms for the CRED-trained BioBERT+SVM model. We restrict our focus to true positives, with the left panel tested on CRED train data, and the right tested on CRED test data. Here p-value is calculated using one-sided unpaired Wilcoxon test as mentioned in Methods, (c) Comparison of importance scores of CRED-trained vs. CDR-trained models for the top 25 words chosen based on the CRED-trained model. (See Figure S17 in the Appendix for similar plot using top 25 words chosen based on the CDR-trained model.), (d) Word cloud of the example abstract shown in Figure S1 based on the importance scores calculated using the CRED-trained (left) and CDR-trained (right) models. Larger word size refers to higher importance-score words focused by the model.

To get further insights into the models, we sort the words based on importance scores and take the top 25 words from CRED- and CDR-trained models separately. In detail, for a given model, we focus on the true positives when the model is applied on the test data (i.e., all test (*G*-*D, Abstract*) tuples that are correctly classified), and then sort the corresponding (*w, G*-*D, Abstract*) tuples based on their importance scores. We can observe from Figure 4c that the top 25 important words of the CRED-trained model contain words like “cause”, and that their importance scores in the CDR-trained model are lower. This trend observed for the BioBERT+SVM models also holds for the BioBERT+XGBoost models (Figures S18 and S19 in Appendix). Although not all 25 words carry a causal connotation, due to contextualized embeddings that consider the entire context, it still emphasizes certain words with causal implications.

We constructed the word cloud of an example abstract from our dataset (the same abstract shown in Figure S1) after training the model on the CRED and CDR datasets separately. From these two word clouds shown in Figure 4d, we can see that the “caused” word is larger in size in the left word cloud as compared to the right one, implying that the CRED-trained model has given the word higher importance. All these model interpretation results taken together demonstrate the benefit of using an expressly-curated causal dataset to train models to extract causal relations.

### Applications of CRED

### CRED-trained model helps predict Parkinson’s disease (PD) genes from PubMed abstracts

To test our CRED-trained BioBERT+SVM model in a real-world setting, we applied our model to all abstracts in PubMed related to Parkinson’s Disease (PD). We downloaded 1,49,945 such abstracts in PubTator format, out of which 45,526 abstracts mentioned at least one gene (see also Methods and Section A.4 of Appendix). Causal predictions for the extracted G-D pairs in these abstracts (specifically 1,11,419 (G-D,Abstract) tuples, with disease being PD) are shown in Figure 5a. We see that many genes (10,919) are less studied, with fewer than 100 papers available to establish their causal relation to disease. However, of the remaining 111 genes, which are present in at least 100 papers, 6 genes are predicted to be causal in at least 50 papers. These six genes were found to be already linked to PD in books^29,30^, which were not part of CRED and hence not used to train our model. Our CRED-trained classifier can thus recapitulate known PD-causing genes from abstracts and quantify the strength of evidence of the causal relation of several genes to a disease like PD.

**Figure 5.**
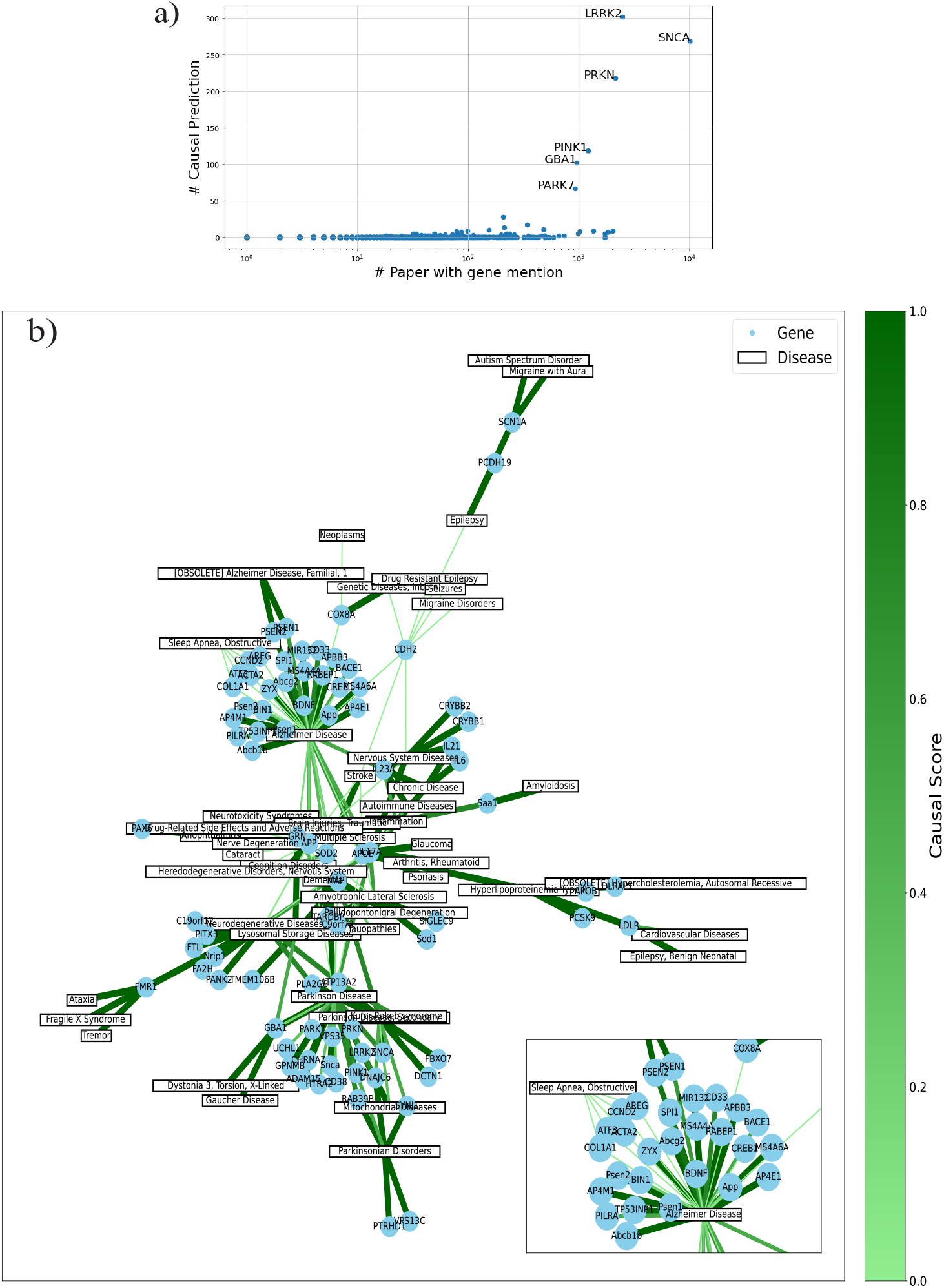
Application results. a) Application of our model to predict PD-causing genes: For each gene, the plot shows the # (number of) PubMed abstracts co-mentioning the gene and PD vs. # abstracts in which our model predicts the gene to be causal for PD. b) Graph of top 250 causal G-D pairs; edge thickness and color are proportional to the CRED-wide causality score. The inset figure is a zoomed-in view of the network centered at the Alzheimer’s disease node. An alternate view focused on the top 20 causal G-D pairs is in Figure S20 in Appendix. Some nodes are mapped to old MeSH IDs (obsolete now) by PubTator – we include them in this figure for completeness.

### CRED-wide causality score for each G-D pair

As a G-D pair can be mentioned in multiple PubMed abstracts, and each such abstract can support the pair as either causal or non-causal, we have a need to derive a dataset-wide causality score of a G-D pair. Focusing on all the PubMed abstracts annotated in CRED, we simply take the ratio of the number of abstracts where a G-D pair is annotated as causal to the number of all abstracts mentioning the G-D pair to obtain a single aggregate causality score for the G-D pair.

Figure 5b shows the top 250 G-D pairs based on the causality score from CRED. In this graph, we can observe causal genes that are common across diseases. For instance, *SCN1A* is causal for Epilepsy, Autism Spectrum Disorder, and Migraine with Aura; and *FMR1* is causal for multiple diseases like Tremors and Ataxia with high causality score. CRED can thus be helpful in determining the causal genes of not only a particular disease, but also common genes that cause different diseases. Such genes can aid target drug discovery efforts pertaining to multiple diseases.

## Discussion

This work proposes CRED, an intra/inter-sentence causal relation extraction (CRE) dataset between genes and diseases from biomedical literature. This work is motivated by the scarcity of annotated biomedical CRE datasets, which have hindered the development of a causal text mining system. Our benchmark CRED (before augmentation) covers 267 abstracts and 3639 gene-disease pairs, out of which 397 are causal; and can be used to train a classification model to perform CRE from text. While the dataset is moderate-sized, its quality compensates for quantity, offering a robust starting point in a field where such annotated datasets spanning multiple genes and diseases are not publicly available. Its manual annotations provide a reference for validating automated tools, benefiting researchers across the biomedical NLP community.

Comparison of our classification model, BioBERT+SVM, with other SOTA models, including LLMs, on our benchmark dataset establishes the unique value our CRED dataset brings to predict causal gene-disease relations. Interpretations of model predictions show that our classifier focused on words with causal connotations for making the predictions. In the application of our CRED-trained BioBERT+SVM classifier on all the abstracts on Parkinson’s disease, we observed that genes predicted to be causal in at least 50 papers were found to be already linked to PD in books. We also found less-studied disease-causing genes that could be explored further. In another application, a CRED-wide gene-disease causality score, which aggregates causal evidence across multiple documents, revealed genes that cause different diseases.

To summarize, our CRE Dataset and CRED-trained classifier are novel resources developed expressly for causal text mining; benchmarking and application of these resources yielded promising results. This reflects the high quality of our manually annotated dataset, though medium-sized can serve as a gold standard for training and evaluating automated relation extraction models. This can thus stimulate further research on language models for causal relation extraction, enabling larger-scale studies in the future.

## Acknowledgments

The authors thank members of the Bioinformatics and Integrative Data Science (BIRDS) group at IIT Madras for their valuable feedback during presentations of this work, and Adwait Parsodkar and Ananya Sai for their feedback on the manuscript. This work is supported by the Wellcome Trust/DBT grant IA/I/17/2/503323 awarded to MN, and summer internship offered to AP by Robert Bosch Centre for Data Science and AI at IIT Madras.

## Author contributions statement

NB and MN conceived and formulated this study, including its methodology, analyses and applications. NB performed all key analyses in the manuscript and interpretation of the results with inputs from MN. NB and SDRC annotated the PubMed abstracts in the CRED dataset. AP conducted analyses related to applications of CRED. NB and MN wrote and reviewed the manuscript. MN guided and supervised the study.

## Data and Code Availability

The CRED dataset and associated code are publicly available at https://github.com/BIRDSgroup/CRED/.

