## Supplementary Information for "Beyond associations: A benchmark Causal Relation Extraction Dataset (CRED) of disease-causing genes, its comparative evaluation, interpretation and application"

### Appendices

#### Appendix A: Supplementary Text

##### Appendix A.1: Related Work

This section summarizes existing datasets and NLP (Natural Language Processing) methods closely related to the CRE (Causal Relation Extraction) task. Since only a few CRE studies exist, many related works discussed here pertain actually to RE (Relation Extraction), with an emphasis on biomedical RE.

***Existing Datasets:***

There are a few publicly available datasets pertaining to RE from the biomedical literature (see Table 1 in main text). A popular dataset for CDR (Chemical Disease Relation) extraction is the BC5CDR (BioCreative V CDR) corpus[1]; containing 1500 abstracts, split into three sets, 500 each for training, validation, and testing. Although BC5CDR is large enough to fine-tune a large language model and contains inter-sentence relations, it does not focus on extracting causal relations (since both mechanistic causal and biomarker correlative type relations between chemicals and diseases are present in this dataset, with no clear labeling of the type of each relation). RE datasets containing gene-disease (G-D) associations (but with no specific mention of causal vs. non-causal relations) include GAD (Gene Disease Association)[2], NHGRI-EBI GWAS (Genome Wide Association Studies) Catalog[3], and DisGeNET[4]. Other RE datasets include EU-ADR (European Union Adverse Drug Reactions) corpus containing pairwise inter-relationships among drugs, disorders and targets, extracted and annotated from medical case reports (including suspected adverse reactions of drugs[5] which precludes statements of causality), and BioRED[6]. BioRED (Biomedical Relation Extraction Dataset)[6] considers relations between various biological entities like chemical, gene, disease, and cell-line; a small fraction (2%) of its relations are causal gene-disease relations as per annotation guidelines (but authors label it as Positive/Negative Correlations, and call the dataset as RE in their paper).

Moving to CRE, dataset like ADE (Adverse Drug Effects)[7] was created to extract drug-related adverse effects from medical reports. Although it is a causal relation extraction dataset, it only talks about the effects of the drugs from 1470 medical case reports. SemEval (Semantic Evaluation)[8] corpus was built to extract causal relations from general natural language and is not specific to biomedical literature. A CRE dataset pertaining to Sjögren's Syndrome (SS)[9] captures the factors that cause a single disease (SS). As it is not publicly available, we requested the dataset from the authors; and observed that it captures causal relations within a sentence, and not across sentences. It is crucial to extend such efforts to multiple diseases, and to consider inter-sentence or within-paragraph relations, so as to enable more comprehensive causal text mining.

***Existing Methods:***

Several model architectures have been proposed for information extraction from biomedical texts. One of the state-of-the-art models, BioBERT[10], based on the BERT model[11] architecture, has been trained using extensive biomedical literature data and fine-tuned for three downstream tasks, namely, Named Entity Recognition (NER), Relation Extraction (RE), and Question Answering (QA). Other models include SciBERT,[12], BlueBERT[13] and PubMedBERT[14], which were trained on a large corpus pertaining to PubMed; and knowledge-graph based K-BERT[15] illustrated for NER and QA. For example, PubMedBERT, also based on BERT, was trained on a large corpus of biomedical research papers, including abstracts from PubMed and full-text articles from PubMed Central. This corpus contains over 100 million words, making it one of the largest pre-training corpora for biomedical text. An inter-sentence RE model[16] is based on a graph convolutional neural network, which is designed to capture local and non-local (e.g., intra- and inter-sentence) dependencies in a document. This model has good performance for RE, but its applicability to CRE was not discussed. Another inter-sentence RE model is BERT-GT[17], and it is focused on extracting relations between chemicals and diseases from the CDR dataset. The SS CRE model[9] leverages causal relations expressed in text using phrases like "leads-to" and "causes"; but this model is limited to the single disease SS and to CRE extraction within sentences. A survey paper[18] reviews works on biomedical causal text mining, and highlights the lack of a benchmark dataset for biomedical CRE.

##### Appendix A.2: Fine-tuning process and parameter calculation

***Fine-tuning BioBERT:***

For fine-tuning BioBERT classifier on the whole abstract embeddings (of length $512 \times 768$), we fine-tune only the parameters of a single hidden-layer neural network classifier that comprises 512 hidden neurons, each of which is connected to 768 neurons of BioBERT's output embedding layer. Embedding of each token (768 dimensions) is reduced to 1 dimension using average pooling, and 512 elements are passed to the output layer. The output from the hidden layer (512 elements) is transformed into the one final output (for binary classification). All parameters of the base BioBERT model are frozen. The total number of parameters (weights and biases) in this fine-tuned neural network classifier stacked at the end of the base BioBERT model can then be calculated as follows:

From the input to the hidden layer: 768 (input features) × 512 (neurons) weights + 512 biases = 393,728 parameters.
From the hidden to the output layer (a sigmoid function): 512 (neurons in hidden layer) weights + 1 bias = 513 parameters.
Total parameters = 393,728 (hidden layer) + 513 (output layer) = 394,241 parameters.
So, the total parameters of Fine-tuned BioBERT classifier is 110 M + 394,241, out of which we trained 394,241 parameters during the fine-tuning process.

***BioBERT + SVM:***
Total number of parameters of BioBERT+SVM is 110 M (base BioBERT) + 4541 (total 4541 training data points leading to 4541 parameters in the Dual SVM problem, with 711 of them being non-zero and corresponding to support vectors). We have reported the difference in the number of parameters of both models in Table S3 in Appendix.
For training of the SVM classifier, the embeddings are of total dimension 4541×1536, where 4541 is the number of training data points, and 1536 = 2×768 is the concatenation of the 768-long embedding of the gene mention and that of the disease mention in the abstract.

#### Appendix A.3: Model Interpretation Details

The importance score formula seen in Methods in main text contains a $(p_{orig} - p_{pert})$ term concerning the prediction probabilities output by the classification model. For SVM classifiers, since it is not based on a probabilistic model, (i) the decision function scores that capture distance from the decision boundary is used to determine $predict_{orig}$ and $predict_{pert}$; and (ii) the probabilities required for calculating $(p_{orig} - p_{pert})$ term in the importance score are obtained using standard methods (Platt scaling as implemented in Python scikit library mentioned in Table S2).

In certain analyses, we need to aggregate the importance score (contribution) of a token $w$ in the $Abstract$ towards all output predictions into a single value, and we achieve it as follows:

$$ImpScore(w, Abstract) = \max_{\text{all G}-\text{D pairs in Abstract}} ImpScore(w, G{-}D, Abstract) \quad \text{(S1)}$$

We selected maximum as the aggregate function in Equation S1, because considering that the same token $w$ can have different importance scores towards different $G{-}D$ pairs in the $Abstract$, we wanted to value a word as important even if it contributes to the causal prediction of just one pair.

In model interpretation analyses, whenever we compare the importance scores of words corresponding to synonyms of "causal" vs. "associate" based on a CRED-trained or CDR-trained model, we employ the unpaired Wilcoxon test that is one-sided (with alternative hypothesis being causal-connotation words having higher importance scores than associate-related words) to obtain the corresponding reported P-value.

#### Appendix A.4: Applications of CRED - Methodological Details

***Predicting the causal genes of Parkinson's disease using all relevant papers in PubMed:***
To apply our model to real-world data, we selected Parkinson's disease (PD). We searched published articles related to PD from PubMed[19] using the keyword "parkinson's", which were 1,49,945 (as of 11-07-2024). We extracted them in Pubtator format (as of 11-07-2024), recognizing all the genes in the abstracts (only 45526 abstracts contained genes). Then, we predict all extracted G-D pairs (1,11,419) (here, the disease is only Parkinson's), using our CRED-trained BioBERT+SVM model, to find all the causal genes of Parkinson's disease in the biomedical literature.

***CRED-wide causality score of a G-D pair:***
To obtain a single causality score of a G-D pair across multiple annotated documents (PubMed abstracts) in CRED, we calculate the ratio of the number of abstracts where the G-D pair is annotated as causal in CRED to the number of abstracts where the G-D pair is mentioned in CRED.

#### Appendix A.5: Synonyms

Synonyms of cause used are: 'causation', 'do', 'inducer', 'induction', 'make', 'inducement', 'stimulate', 'stimulus', 'inducive', 'get', 'causative', 'have', 'cause', 'induce', 'stimulation', 'inducing', 'causation', 'impact', 'developement', 'get', 'make', 'case', 'movement', 'induce', 'stimulus', 'causal', 'crusade', 'do', 'inducer', 'inducement', 'causal-agent', 'campaign', 'causative', 'grounds', 'reason', 'stimulation', 'drive', 'stimulate', 'cause', 'induction', 'causa', 'inducive', 'effort', 'suit', 'inducing', 'caused', 'causes'.

Synonyms of associate used are: 'associate', 'associations', 'connective', 'associatory', 'fellow', 'tie-in', 'connexion', 'associate-degree', 'associable', 'familiar', 'affiliate', 'companion', 'fellowship', 'link', 'linked', 'support', 'related', 'consort', 'colligation', 'relation', 'association', 'comrade', 'connect', 'associative', 'linkage', 'connection', 'link-up', 'correlate', 'correlative', 'correlativity', 'correlated', 'correlation'.

#### Appendix B: Supplementary Figures

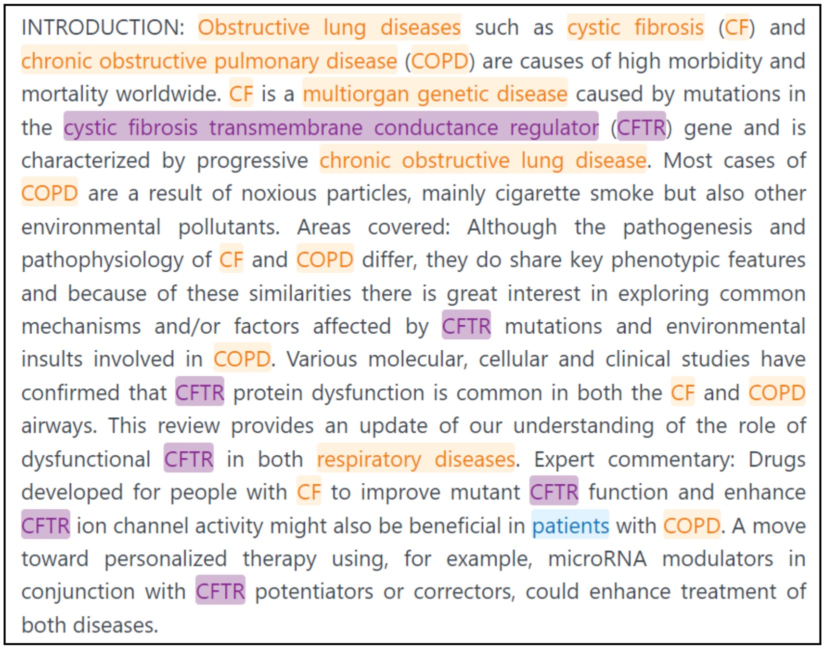

INTRODUCTION: Obstructive lung diseases such as cystic fibrosis (CF) and chronic obstructive pulmonary disease (COPD) are causes of high morbidity and mortality worldwide. CF is a multiorgan genetic disease caused by mutations in the cystic fibrosis transmembrane conductance regulator (CFTR) gene and is characterized by progressive chronic obstructive lung disease. Most cases of COPD are a result of noxious particles, mainly cigarette smoke but also other environmental pollutants. Areas covered: Although the pathogenesis and pathophysiology of CF and COPD differ, they do share key phenotypic features and because of these similarities there is great interest in exploring common mechanisms and/or factors affected by CFTR mutations and environmental insults involved in COPD. Various molecular, cellular and clinical studies have confirmed that CFTR protein dysfunction is common in both the CF and COPD airways. This review provides an update of our understanding of the role of dysfunctional CFTR in both respiratory diseases. Expert commentary: Drugs developed for people with CF to improve mutant CFTR function and enhance CFTR ion channel activity might also be beneficial in patients with COPD. A move toward personalized therapy using, for example, microRNA modulators in conjunction with CFTR potentiators or correctors, could enhance treatment of both diseases.

**Figure S1.** Example of an abstract (PMID: 29750581) from our dataset taken from Pubtator[14]

.

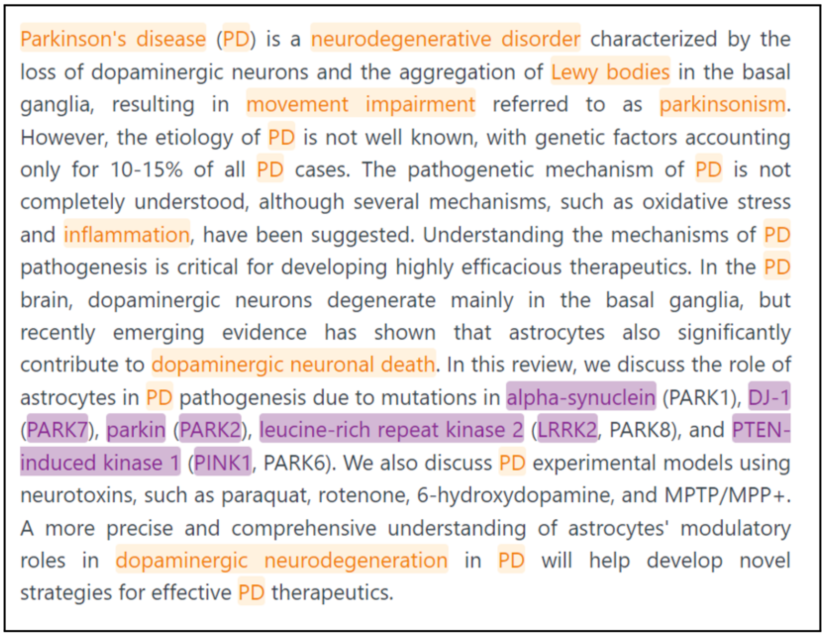

Parkinson's disease (PD) is a neurodegenerative disorder characterized by the loss of dopaminergic neurons and the aggregation of Lewy bodies in the basal ganglia, resulting in movement impairment referred to as parkinsonism. However, the etiology of PD is not well known, with genetic factors accounting only for 10-15% of all PD cases. The pathogenetic mechanism of PD is not completely understood, although several mechanisms, such as oxidative stress and inflammation, have been suggested. Understanding the mechanisms of PD pathogenesis is critical for developing highly efficacious therapeutics. In the PD brain, dopaminergic neurons degenerate mainly in the basal ganglia, but recently emerging evidence has shown that astrocytes also significantly contribute to dopaminergic neuronal death. In this review, we discuss the role of astrocytes in PD pathogenesis due to mutations in alpha-synuclein (PARK1), DJ-1 (PARK7), parkin (PARK2), leucine-rich repeat kinase 2 (LRRK2, PARK8), and PTEN-induced kinase 1 (PINK1, PARK6). We also discuss PD experimental models using neurotoxins, such as paraquat, rotenone, 6-hydroxydopamine, and MPTP/MPP+. A more precise and comprehensive understanding of astrocytes' modulatory roles in dopaminergic neurodegeneration in PD will help develop novel strategies for effective PD therapeutics.

**Figure S2.** Annotation discrepancy example PMID (36831289) For alpha-synuclein → PD: Annotator 1: Non-causal and Annotator 2: causal

Parkinson's disease (PD) is defined as a complex disorder with multifactorial pathogenesis, yet a more accurate definition could be that PD is not a single entity, but rather a mixture of different diseases with similar phenotypes. Attempts to classify subtypes of PD have been made based on clinical phenotypes or biomarkers. However, the most practical approach, at least for a portion of the patients, could be to classify patients based on genes involved in PD. GBA and LRRK2 mutations are the most common genetic causes or risk factors of PD, and PRKN is the most common cause of autosomal recessive form of PD. Patients carrying variants in GBA, LRRK2 or PRKN differ in some of their clinical characteristics, pathology and biochemical parameters. Thus, these three PD-associated genes are of special interest for drug development. Existing therapeutic approaches in PD are strictly symptomatic, as numerous clinical trials aimed at modifying PD progression or providing neuroprotection have failed over the last few decades. The lack of precision medicine approach in most of these trials could be one of the reasons why they were not successful. In the current review we discuss novel therapeutic approaches targeting GBA, LRRK2 and PRKN and discuss different aspects related to these genes and clinical trials.

**Figure S3.** Annotation discrepancy example PMID (34626666) For GBA → PD and LRRK2 → PD: : Annotator 1: Causal and Annotator 2: Non-causal

Original abstract:
Genetic studies have led to major discoveries in the pathogenesis of various neurodegenerative diseases. Ubiquitin-positive familial frontotemporal dementia was recently found to be caused by mutations in the Progranulin gene (PGRN), and the major constituent of the inclusions, TDP-43, was subsequently identified.

Augmented abstract:
genetic studies have shown the important discoveries in the pathogenesis of different neurodegenerative diseases. ubiquitin - positive familial frontotemporal dementia was discovered in be caused by mutations in the progranulin gene (pgrn), where main constituent of the inclusions, tdp - 43, was subsequently recognized.

**Figure S4.** Example showing original and its augmented abstract (PMID:18322368) from CRED

| Hyper Para | Values | Best |
|---|---|---|
| Kernel | RBF, Linear, Poly | Poly |
| C | 0.1, 1, 10 | 10 |
| Degree | 6, 10, 20, 30 | 10 |
| Gamma | Scale, Auto | Scale |
| Class Weight | 0:1 1:10, 0:1 1:30, 0:1 1:50 | 0:1 1:30 |

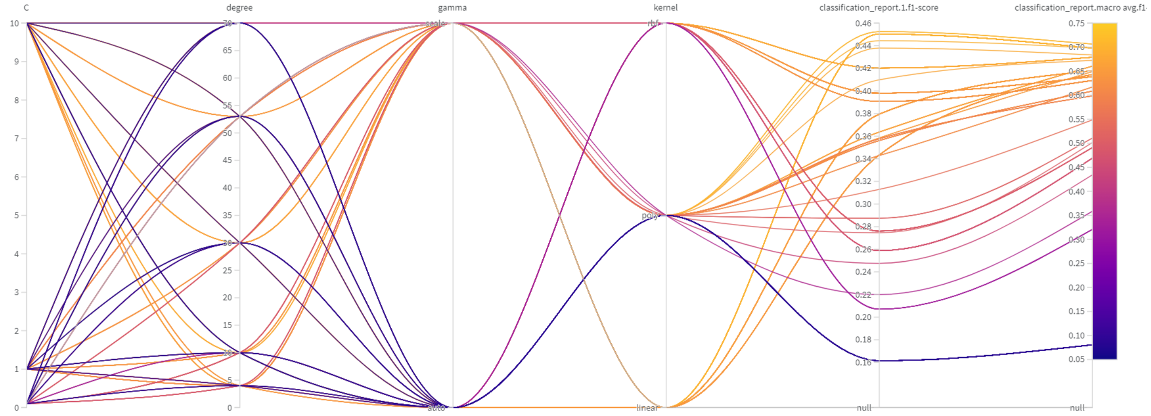


**Figure S5.** Hyper parameters tuning for BioBERT+SVM

| Hyper Parameters | Values | Best |
|---|---|---|
| Learning Rate | 0.01, 0.1, 0.2 | 0.1 |
| Max Depth | 5, 10, 20, 40 | 40 |
| No. of estimators | 100, 200, 400 | 400 |
| Gamma | 0.1, 0.2, 0.3 | 0.1 |
| Scale pos weight | 100, 200, 500 | 500 |

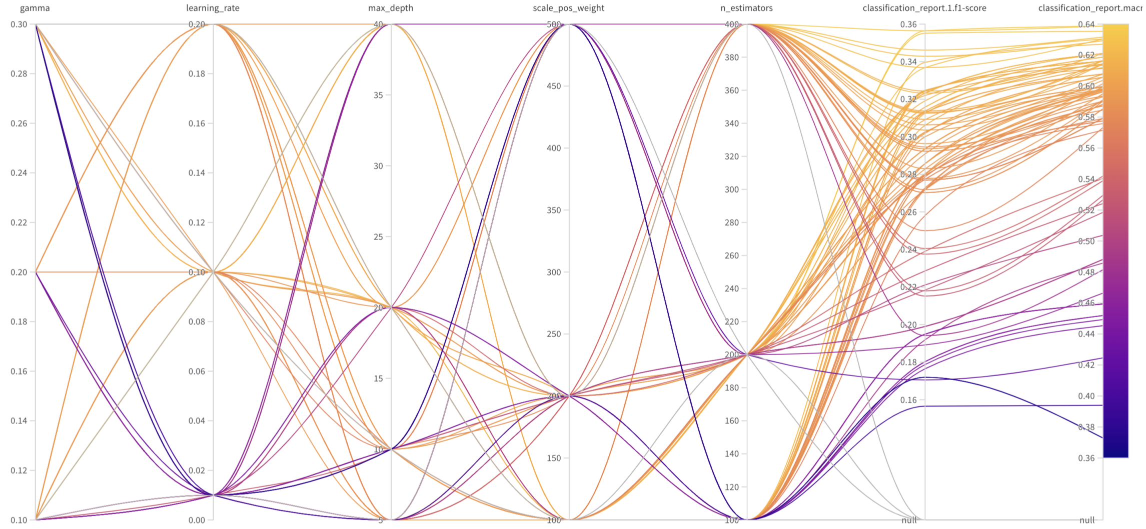


**Figure S6.** Hyper parameters tuning for BioBERT+XG Boost

| Hyper Parameters | Values | Best |
|---|---|---|
| No. of jobs | 2,5,10 | 5 |
| Max Depth | 10, 50 , 100 | 50 |
| No. of estimators | 100, 200, 400 | 200 |
| Max leaf nodes | 100, 200, 500 | 500 |
| Max features | Auto, Sqrt | Sqrt |
| Class Weight | 0:1 1:10, 0:1 1:30, 0:1 1:50 | 0:1 1:30 |

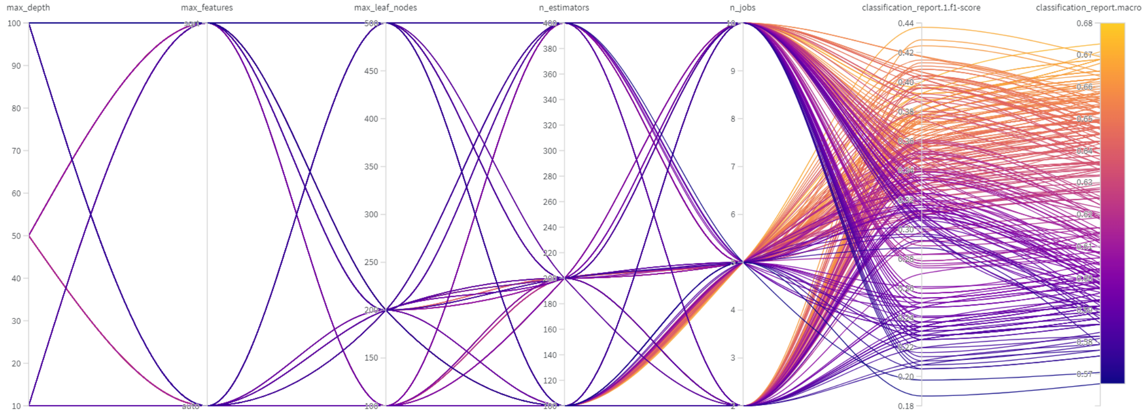


**Figure S7.** Hyper parameters tuning for BioBERT+Random Forest

| Hyper Para | Values | Best |
|---|---|---|
| Learning Rate | 2e-5, 3e-5, 5e-5 | 3e-5 |
| No. of train epochs | 1, 2, 3, 4 | 4 |
| Weight decay | 0.01, 0.05, 0.1 | 0.05 |
| Warmup Steps | 0, 500, 1000 | 1000 |

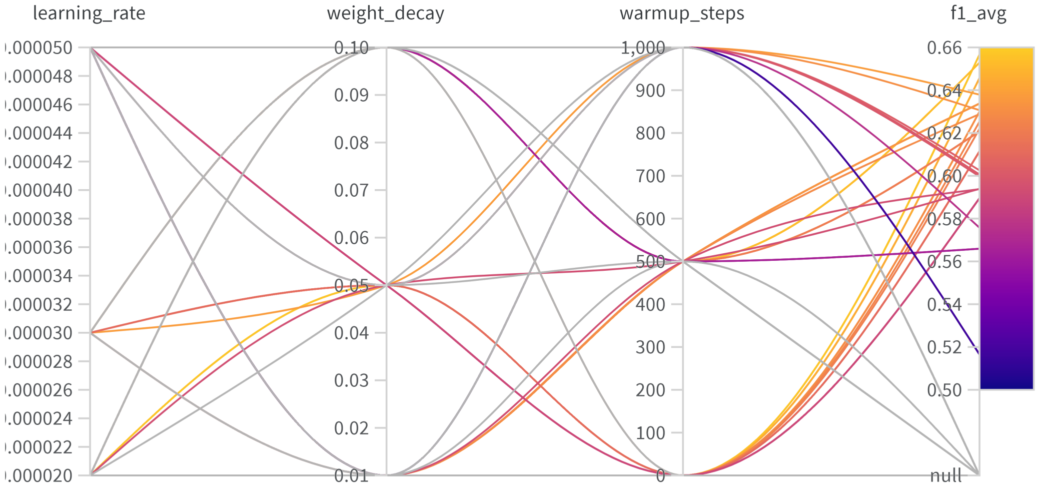


**Figure S8.** Hyper parameters tuning for BioBERT

| Hyper Para | Values | Best |
|---|---|---|
| Learning Rate | ’2e-5, 3e-5, 5e-5 | 2e-5 |
| No. of train epochs | 1, 2, 3, 4 | 3 |
| Weight decay | 0.01, 0.05, 0.1 | 0.05 |
| Warmup steps | 0, 500, 1000 | 0 |

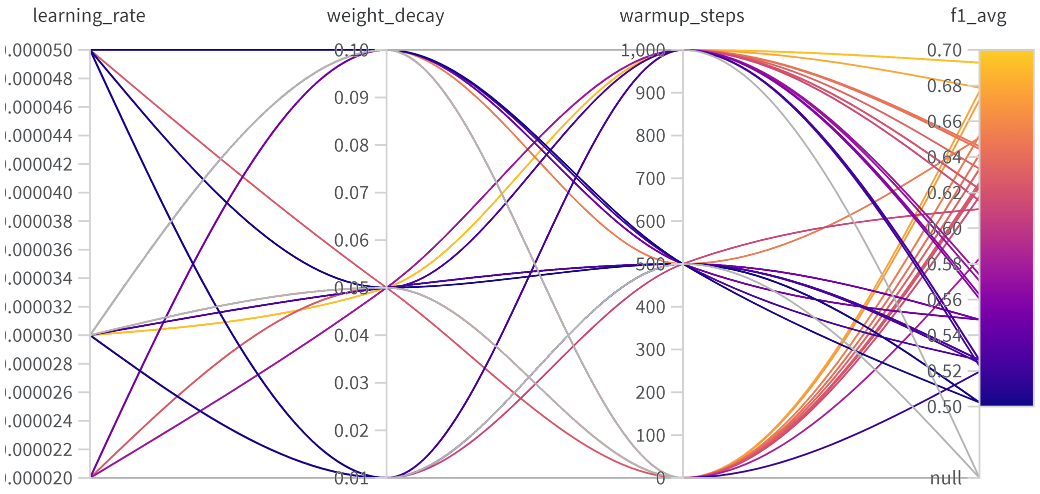


**Figure S9.** Hyper parameters tuning for PubMedBERT

| Hyper Para | Values | Best |
|---|---|---|
| Learning Rate | ’2e-5, 3e-5, 5e-5 | 3e-5 |
| No. of train epochs | 1, 2, 3, 4 | 2 |
| Weight decay | 0.01, 0.05, 0.1 | 0.05 |
| Warmup steps | 0, 500, 1000 | 0 |

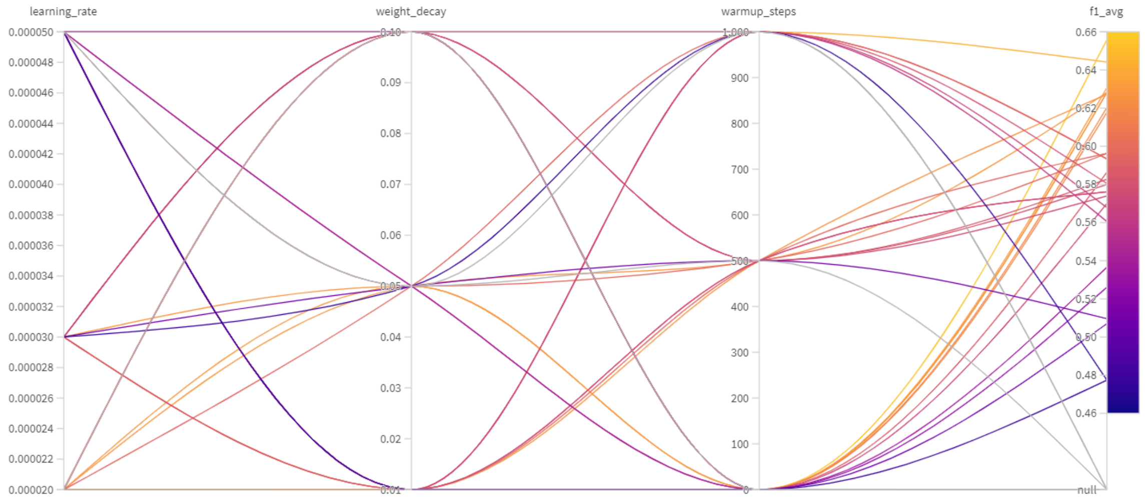


**Figure S10.** Hyper parameters tuning for SciBERT

| Hyper Para | Values | Best |
|---|---|---|
| Learning Rate | ’2e-5, 3e-5, 5e-5 | 3e-5 |
| No. of train epochs | 1, 2, 3, 4 | 3 |
| Weight decay | 0.01, 0.05, 0.1 | 0.05 |
| Warmup steps | 0, 500, 1000 | 0 |

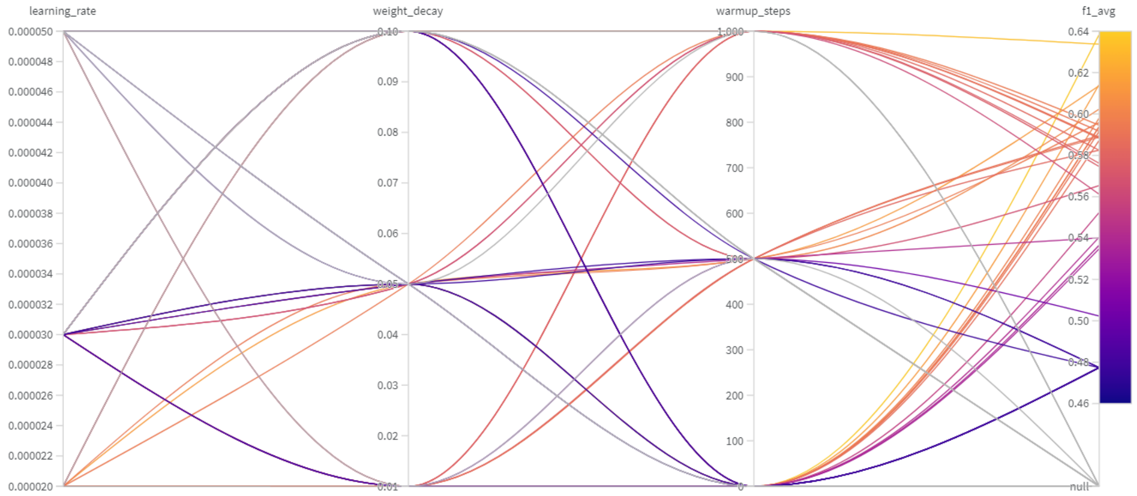


**Figure S11.** Hyper parameters tuning for BlueBERT

| Hyper Para | Values | Best |
|---|---|---|
| Kernel | RBF, Linear, Poly | Linear |
| C | 0.1, 1, 10 | 10 |
| Degree | 10, 30, 50, 70 | 70 |
| Gamma | Scale, Auto | Auto |
| Class Weight | 0:1 1:10, 0:1 1:30, 0:1 1:50 | 0:1 1:30 |

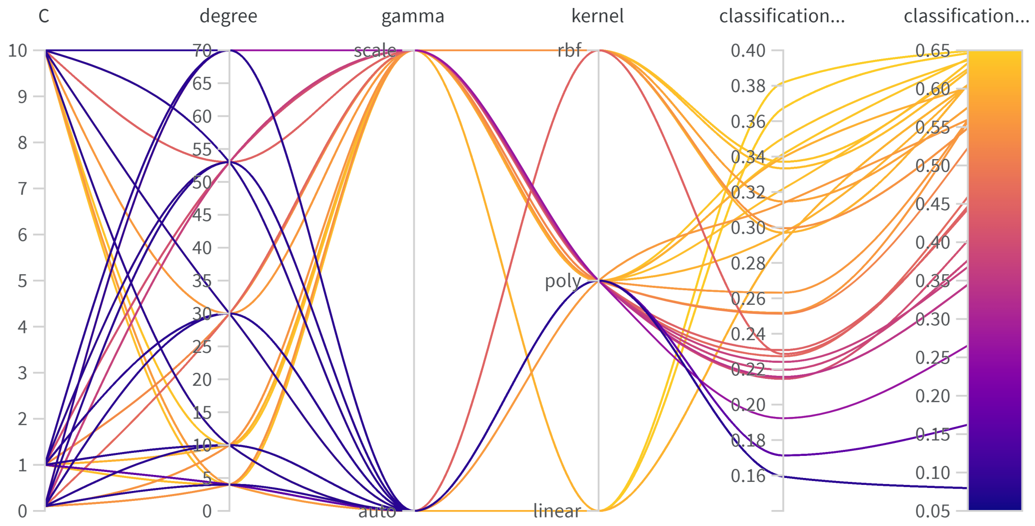


**Figure S12.** Hyper parameters tuning for SciBERT+SVM

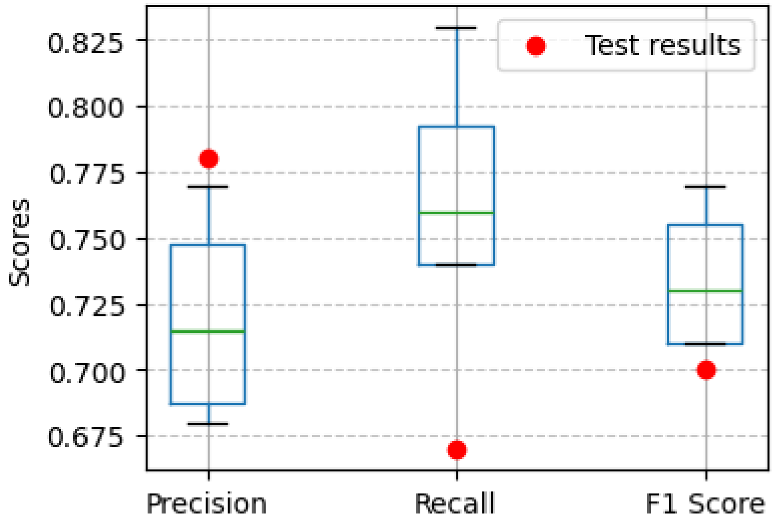


**Figure S13.** Performance of BioBERT+SVM model trained on different three-fold combinations and tested on held-out validation fold and test data.

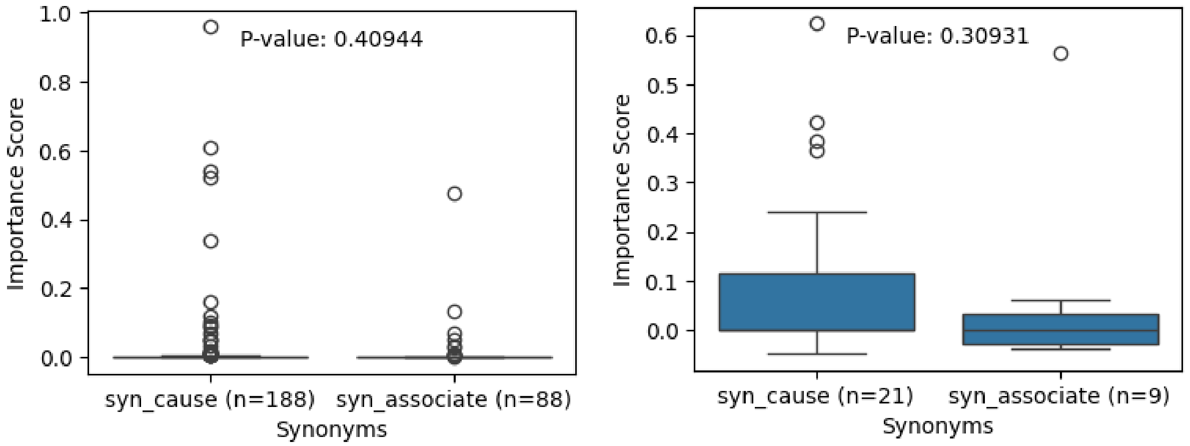


**Figure S14.** Comparison of importance scores of “cause” vs. “associate” synonyms for CRED-trained BioBERT+XGBoost model. We restrict our focus to true positives, with the left panel tested on CRED train data, and the right tested on CRED test data. Here p-value is calculated using one-sided unpaired Wilcoxon test as mentioned in Methods. For this CRED-trained XGBoost model, which has poorer classification performance than the BioBERT+SVM model, the trend of causal words having higher importance scores than associate-related words was weaker and not statistically significant.

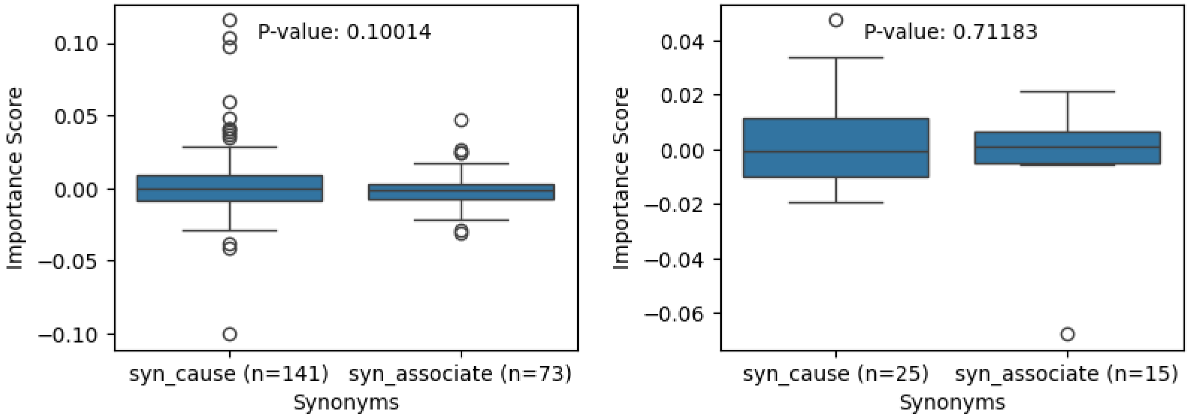


**Figure S15.** Comparison of importance scores of “cause” vs. “associate” synonyms for CDR-trained BioBERT+SVM model. We restrict our focus to true positives, with the left panel tested on CRED train data, and the right tested on CRED test data. Here p-value is calculated using one-sided unpaired Wilcoxon test as mentioned in Methods. In the train data, the P-value is weakly significant (P-value 0.10014). In the test data, which better captures the model behavior than the train data, model does not show higher importance scores of causal over associate synonyms (P-value 0.71). This behavior may be due to the model focusing more on associate than causal synonyms (which is consistent with the model being CDR-trained).

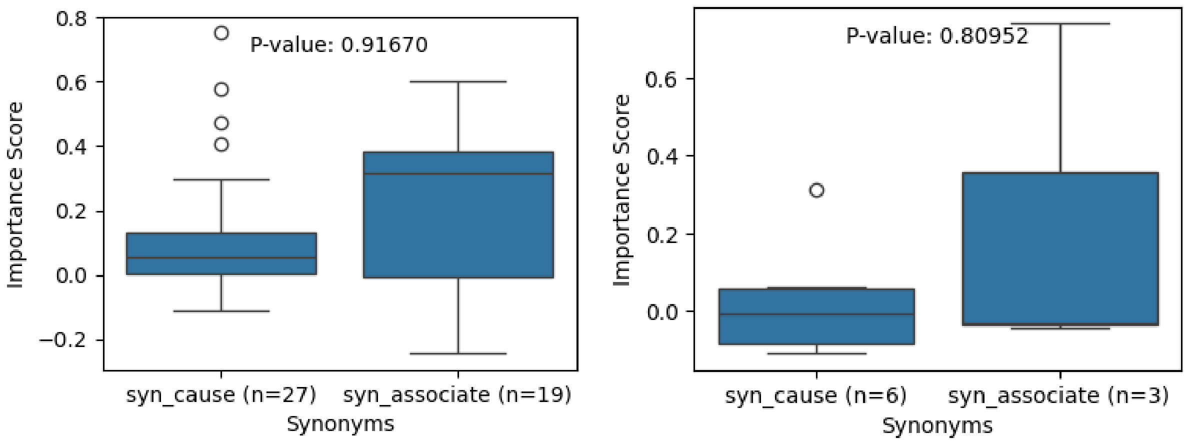


**Figure S16.** Comparison of importance scores of "cause" vs. "associate" synonyms for CDR-trained BioBERT+XGBoost model. We restrict our focus to true positives, with the left panel tested on CRED train data, and the right tested on CRED test data. Here p-value is calculated using one-sided unpaired Wilcoxon test as mentioned in Methods. In the train data, the P-value is not significant (P-value 0.91670). In the test data, which better captures the model behavior than the train data, model does not show higher importance scores of causal over associate synonyms (P-value 0.81). This behavior may be due to the model focusing more on associate than causal synonyms (which is consistent with the model being CDR-trained).

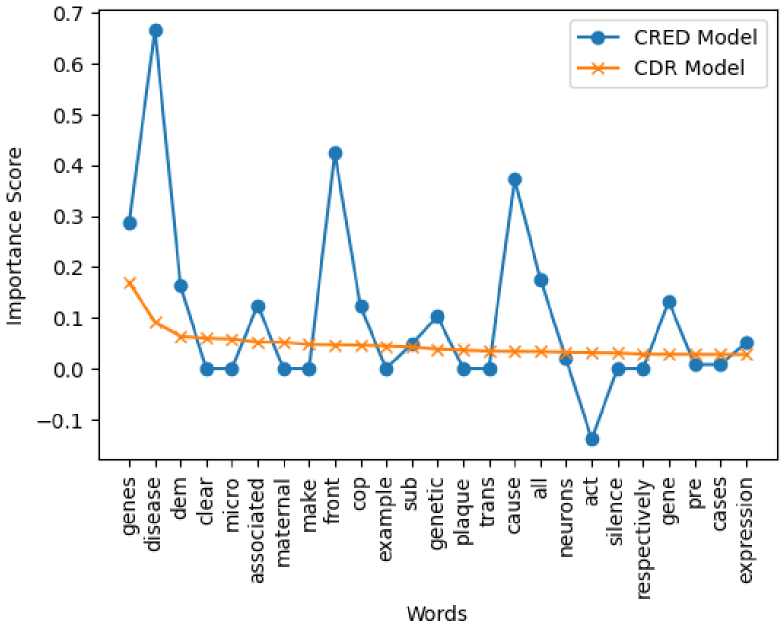


**Figure S17.** Comparison of importance scores of top 25 words (from CDR-trained model) in CRED-trained and CDR-trained SVM models.

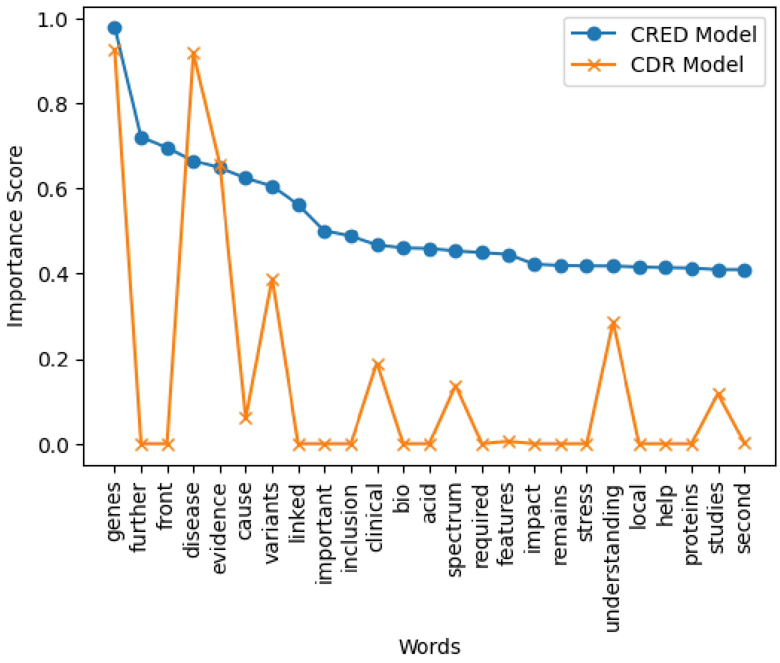


**Figure S18.** Comparison of importance scores of top 25 words (from CRED-trained model) in CRED-trained and CDR-trained XGBoost models.

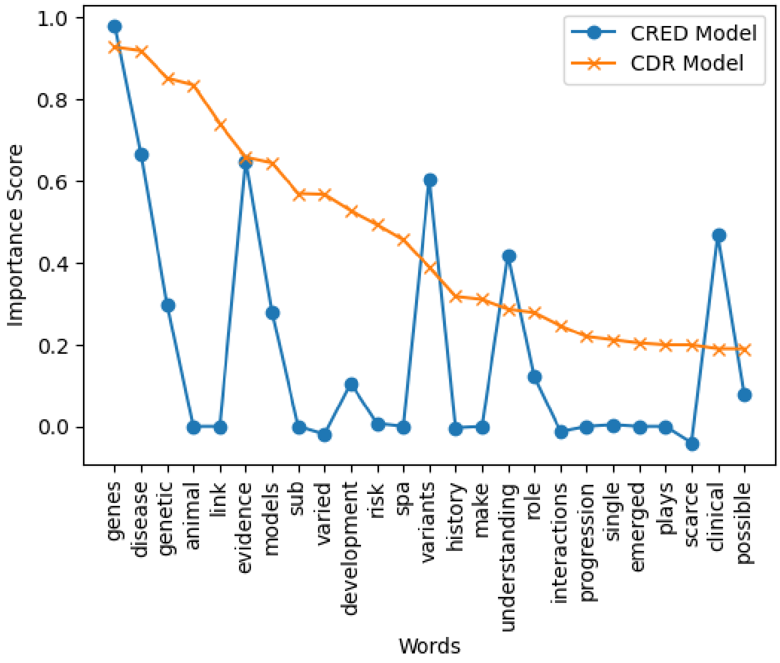


**Figure S19.** Comparison of importance scores of top 25 words (from CDR-trained model) in CRED-trained and CDR-trained XGBoost models.

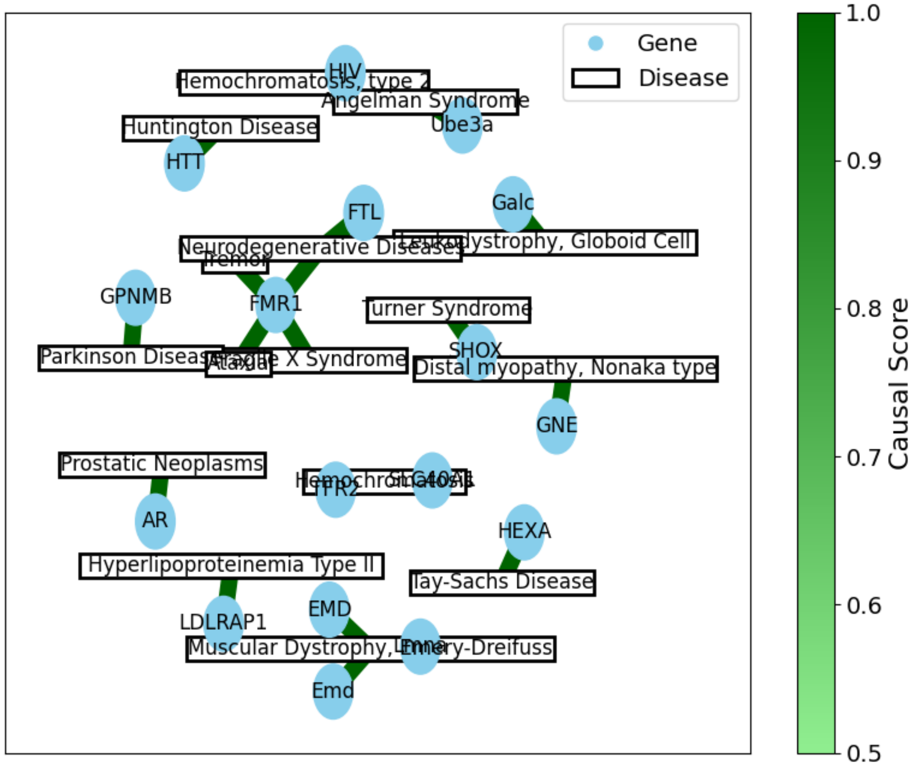


**Figure S20.** Graph of top 20 causal G-D pairs; edge thickness and color shown are proportional to the CRED-wide causality score.

#### Appendix C: Supplementary Tables

**Table S1.** Performance of BioBERT+SVM model trained on CRED training data before augmentation and evaluated on CRED test data.

| | Positive class | | | Average | | |
|---|---|---|---|---|---|---|
| | **Prec** | **Rec** | **F1** | **Prec** | **Rec** | **F1** |
| **SVM** | 0.35 | 0.35 | 0.35 | 0.64 | 0.64 | 0.64 |

**Table S2.** Software/Packages specifications

| Package/Software | Version |
|---|---|
| matplotlib | 3.8.0 |
| nltk | 3.8.1 |
| numpy | 1.26.0 |
| python | 3.10.0 |
| scikit-learn | 1.3.1 |
| scipy | 1.11.3 |
| seaborn | 0.13.0 |
| torch | 2.0.1 |
| tqdm | 4.66.1 |
| wandb | 0.15.11 |
| wordcloud | 1.9.2 |
| xgboost | 2.0.0 |

**Table S3.** Parameters of models. See Appendix A.2 for how these numbers are derived.

| | Total Params | Trained Params |
|---|---|---|
| **BioBERT** | 110 M + 394241 | 394241 |
| **BioBERT + SVM** | 110 M + 711 | 711 |

**Table S4.** Performance of BioBERT+SVM model, when trained on CRED and tested on different datasets. Prec, Rec, and F1 denote precision, recall, and F1 score respectively.

| | Positive class | | | Average | | |
|---|---|---|---|---|---|---|
| **Tr-Te** | **Prec** | **Rec** | **F1** | **Prec** | **Rec** | **F1** |
| **CRED-CDR** | 0.27 | 0.65 | 0.39 | 0.58 | 0.62 | 0.55 |
| **CRED-BioRED** | 0.06 | 0.16 | 0.09 | 0.48 | 0.45 | 0.45 |
| **CRED-GAD** | 0.44 | 0.004 | 0.01 | 0.46 | 0.50 | 0.32 |
